## Supplemental text. for "Evolution of phenotypic plasticity leads to tumor heterogeneity with implications for therapy"

### 1 Model definition

#### 1.1 State space

We define a lattice-gas cellular automaton (LGCA) model on a one-dimensional lattice $\mathcal{L} \subset \mathbb{Z}$. Every lattice node $\mathbf{r} \in \mathcal{L}$ contains two velocity channels $c_i = \pm 1, i = 0, 1$ representing movement to the right and left, respectively. Additionally, one rest channel $c_2 = 0$ represents no movement. We use an LGCA variant called evolutionary LGCA (evo-LGCA), in which there is no exclusion principle, and the presence of cells is represented by properties $p \in \mathcal{P}$. Here, the unique property is the switch regulation $\kappa \in \mathbb{R}$, i. e., $\mathcal{P} = \mathbb{R}$. A finite multiset of cell properties indicates the occupation of a channel (channel occupation). A multiset $M$ is a set that can contain the same element multiple times, i. e., $M : \mathcal{P} \mapsto \mathbb{N}_0$, and $M(p)$ indicates the number of cells with the property $p$ (multiplicity of $p$). Here, we are only interested in finite multisets representing finite cell numbers, i. e., $\operatorname{supp}(M) := \{p \in \mathcal{P} \mid M(p) \neq 0\}$, $|\operatorname{supp}(M)| < \infty$. The set of all possible finite multisets on $\mathcal{P}$ is defined as

$$\mathcal{M}_{\mathcal{P}} := \left\{M \in \mathbb{N}_0^{\mathcal{P}} \mid |\operatorname{supp}(M)| < \infty\right\}, \tag{S1}$$

where $\mathbb{N}_0^{\mathcal{P}}$ denotes the set of all multisets on $\mathcal{P}$. Then, the state space of a node in the evo-LGCA framework is defined as $\mathcal{E} := \mathcal{M}_{\mathcal{P}}^{b+1}$, i. e., every channel contains a finite multiset indicating the presence of cells.

Cells with the migratory phenotype reside in the velocity channels, while cells with the proliferative phenotype are in the rest channel. The state of a node $r$ at time $k$ is given by the configuration $\boldsymbol{\eta}(r,k) = \mathbf{s} \in \mathcal{E}$. The number of cells is given by the sum of the cardinalities of the multisets, i. e.,

$$n(r,k) := \sum_{i=0}^{b} |\eta_i(r,k)|. \tag{S2}$$

**Table S1. List of symbols**

| Symbol | Explanation |
|---|---|
| $\mathcal{L} \subset \mathbb{Z}^d$ | $d$-dimensional regular lattice |
| $b = 2d$ | Number of velocity channels |
| $L \in \mathbb{N}$ | Lattice side length |
| $\mathbf{r} \in \mathcal{L}$ | Lattice node |
| $k \in \mathbb{N}$ | Automaton time step |
| $K > 0$ | Node capacity |
| $n(\mathbf{r}, k) \in \mathbb{N}_0$ | Cell number at node $\mathbf{r}$ and time step $k$ |
| $\rho_{\mathcal{N}}(\mathbf{r}, k) > 0$ | Average density around node $\mathbf{r}$ at time step $k$ |
| $\delta \in [0, 1]$ | Death probability |
| $\alpha(1 - n(\mathbf{r}, k)/K)$ | Proliferation probability of resting cells |
| $r_\kappa \in [0, 1]$ | Probability of switching to the proliferative phenotype |
| $1 - r_\kappa \in [0, 1]$ | Probability of switching to the migratory phenotype |
| $\kappa \in \mathbb{R}$ | Individual phenotypic switch regulation |
| $\theta \in [0, 1]$ | Critical cell density where phenotypes are equally probable |
| $\Delta\kappa \in \mathbb{R}^+$ | Standard deviation of $\kappa$-values of offspring versus mother cells |

**Table S2. Parameter values used in parameter scan**

| Parameter | Value(s) |
|---|---|
| $b$ | 2 |
| $L$ | 1001 |
| $K$ | 100 |
| $\Delta\kappa$ | 0.2 |
| $\alpha$ | 1.0 |
| $\theta$ | $[0.0, 1.0]$ |
| $\delta$ | $[0.0, 0.25]$ |

#### 1.2 Dynamics

##### 1.2.1 Overview

The model dynamics is defined via two operators: a deterministic transport operator $\mathcal{T}$, and a stochastic rearrangement operator $\mathcal{R}$. The latter comprises births and deaths, phenotypic switches, and changes in migration velocity. The transport operator propagates cells in velocity channels towards their respective direction of movement. The composition of the two operators $\mathcal{T} \circ \mathcal{R}$ is applied independently at every node $r$ and at each time step $k$ to create the configuration at the next time step $k+1$.

The stochastic rearrangement operator is defined as a composition of four processes: cell death $\mathcal{D}$, phenotypic switching $\mathcal{S}$, proliferation $\mathcal{B}$, and orientation $\mathcal{O}$,

$$\boldsymbol{\eta} \xrightarrow{\mathcal{D}} \boldsymbol{\eta}' \xrightarrow{\mathcal{S}} \boldsymbol{\eta}'' \xrightarrow{\mathcal{B}} \boldsymbol{\eta}''' \xrightarrow{\mathcal{O}} \boldsymbol{\eta}^{\mathcal{R}}. \tag{S3}$$

The precise updated rules are defined below. Note that we assume that each cell dies, switches phenotype, proliferates, and switches direction of movement independently.

##### 1.2.2 Cell death

We assume that each tumor cell dies independently from other cells with probability $\delta$ in each time step. Therefore, the number of dying cells with property $p$ follows a binomial distribution, and we can write the new configuration after cell deaths as

$$\boldsymbol{\eta}' = \boldsymbol{\eta} - \boldsymbol{\eta}_D, \tag{S4}$$

where $\boldsymbol{\eta}_D$ is a configuration that contains each dying cell.

The sum of channel occupations $a+b$, where $a, b \in \mathcal{M}_{\mathcal{P}}$, is the multiset $c \in \mathcal{M}_{\mathcal{P}}$ defined by its multiplicity

$$M_c(p) = M_a(p) + M_b(p) \quad \forall p \in \mathcal{P}. \tag{S5}$$

Similarly, the difference of two channel occupations $a-b$, where $a, b \in \mathcal{M}_{\mathcal{P}}$, as $c \in \mathcal{M}_{\mathcal{P}}$ is defined as via the multiplicity as

$$M_c(p) = \max(M_a(p) - M_b(p), 0) \quad \forall p \in \mathcal{P}. \tag{S6}$$

The definitions of sum and difference of channel occupations extend to configurations of nodes by point-wise application, i. e., $\mathbf{c} = \mathbf{a} \pm \mathbf{b}$, with $\mathbf{a}, \mathbf{b}, \mathbf{c} \in \mathcal{E}$, is defined as $\mathbf{c} := (a_0 \pm b_0, a_1 \pm b_1, \ldots, a_b \pm b_b)^T$.

$\boldsymbol{\eta}_D$, the node configuration indicating dying cells, is formally defined by the probability

$$P_D(\boldsymbol{\eta}_D|\boldsymbol{\eta}) = \prod_{i=0}^{b} \prod_{\kappa \in \mathrm{supp}(M_{\eta_i})} \binom{M_{\eta_i}(\kappa)}{M_{\eta_{i,D}}(\kappa)} \delta^{M_{\eta_{i,D}}(\kappa)} (1-\delta)^{M_{\eta_i}(\kappa) - M_{\eta_{i,D}}(\kappa)}, \tag{S7}$$

where $M_{\eta_i}(p)$ indicates the number of cells with property $p$ in channel $i$ of node configuration $\boldsymbol{\eta}$.

##### 1.2.3 Phenotypic switch

After the death operator has removed the dying cells, cells switch their phenotype according to the phenotypic switch probability $r_\kappa(\rho_\mathcal{N}) \in [0,1]$ depending on the individual switch parameter $\kappa$ and the average local density $\rho_\mathcal{N}(\mathbf{r},k) := \frac{1}{(b+1)K}\sum_{i=0}^{b} n(\mathbf{r}+\mathbf{c}_i,k)$:

$$r_\kappa(\rho_\mathcal{N}) := \frac{1}{2}(1+\tanh(\kappa(\rho_\mathcal{N}-\theta))). \tag{S8}$$

To express this process mathematically, we first define the multiset of all cells on a node

$$\eta_{\text{tot}}(\mathbf{r},k) := \sum_{i=0}^{b} \eta_i(\mathbf{r},k). \tag{S9}$$

Based on this, we construct the multisets of cells with the migratory $\eta_m$ and proliferative $\eta_p$ phenotypes after the phenotype switch, respectively

$$P_p(\eta''_p|\eta'_{\text{tot}}) = \prod_{\kappa\in\text{supp}\left(M_{\eta'_{\text{tot}}}\right)} \binom{M_{\eta'_{\text{tot}}}(\kappa)}{M_{\eta''_p}(\kappa)} r_\kappa^{M_{\eta''_p}(\kappa)} (1-r_\kappa)^{M_{\eta'_{\text{tot}}}(\kappa)-M_{\eta''_p}(\kappa)} \tag{S10}$$

and

$$\eta''_m := \eta'_{\text{tot}} - \eta''_p. \tag{S11}$$

##### 1.2.4 Proliferation

Cells in the proliferative phenotype produce offspring with probability $P_B = \alpha(1-n/K)$. The offspring is initially in the proliferative phenotype and contained in the offspring multiset $\eta_B$. Each daughter cell's phenotypic switch parameter $\kappa'$ is drawn from a Gaussian probability distribution $f(\kappa'|\kappa)$ centered around their mother cell's phenotypic switch parameter $\kappa$

$$\kappa' \sim \mathcal{N}(\kappa, \Delta\kappa). \tag{S12}$$

The multiset of cells in the proliferative phenotype after the birth step is then

$$\eta'''_p = \eta''_p + \eta_B. \tag{S13}$$

The offspring multiset $\eta_B$ is informally defined as a stochastic process in the following way

$$\eta_B = \left\{\text{Draw } \kappa' \quad \text{from} \quad f(\kappa'|\kappa) \quad \text{with probability} \quad \alpha(1-n/K) \quad \forall\kappa\in\eta''_p\right\}. \tag{S14}$$

##### 1.2.5 Reorientation

In the final substep of the stochastic rearrangement, cells with the migratory phenotype choose their new direction of movement independently and with uniform probability $T_i = 1/b$. Consequently, the probability for the post-orientation configuration $\boldsymbol{\eta}^{\mathcal{R}}$ is a multinomial distribution defined by the transition probability

$$P(\boldsymbol{\eta}^{\mathcal{R}}|\eta'''_m) = \prod_{\kappa\in\mathrm{supp}(\eta'''_m)} M_{\eta'''_m}(\kappa)! \prod_{i=0}^{b-1} \frac{T_i^{M_{\eta_i^{\mathcal{R}}}(\kappa)}}{M_{\eta_i^{\mathcal{R}}}(\kappa)!} \boldsymbol{\delta}\Big(M_{\eta'''_m}(\kappa), M_{\eta^{\mathcal{R}}_{\mathrm{tot}}}(\kappa)\Big). \tag{S15}$$

Here $\boldsymbol{\delta}(x,y) := \begin{cases} 1 & \text{if} \quad x = y, \\ 0 & \text{else} \quad , \end{cases}$ is the Kronecker delta, which ensures mass conservation during the orientation step and must not to be confused with the death probability $\delta$. The rest channel occupation is given by the multiset of cells with the proliferative phenotype after the birth step and unchanged by the orientation step, i. e., $\eta_b^{\mathcal{R}} = \eta_p'''$. Therefore the second product in Eq. (S15) is only multiplying the terms corresponding to the velocity channels (the index runs from $i = 0$ to $i = b - 1$).

#### 1.3 Transport

After stochastic rearrangement, migratory cells move in their direction of movement, while proliferative cells stay in the rest channel of the same node. In other words

$$\eta_i(\mathbf{r}, k+1) = \eta_i^{\mathcal{R}}(\mathbf{r} - \mathbf{c}_i, k). \tag{S16}$$

### 2 Derivation of per-capita growth rate

Here, we derive the average per-capita growth rate of a cell with switch parameter $\kappa$ in a microenvironment with mean density $\rho_{\mathcal{N}}$. The probability for death is equal to $\delta$, i. e., $P(\Delta n = -1) = \delta$. The probability for proliferation is equal to the probability of survival times the probability of being in the proliferative phenotype times the probability of proliferation, i. e., $P(\Delta n = +1) = (1 - \delta)\alpha(1 - \rho)r_\kappa(\rho_{\mathcal{N}})$. Therefore, the average change is

$$F_\kappa(\rho, \rho_{\mathcal{N}}) := P(\Delta n) = (1-\delta)\alpha(1-\rho)r_\kappa(\rho_{\mathcal{N}}) - \delta \approx \alpha(1-\rho)r_\kappa(\rho_{\mathcal{N}}) - \delta \quad \text{for} \quad \delta \ll 1. \tag{S17}$$

### 3 Scaling of LGCA dimensions

#### 3.1 Maximum tumor size

Here, we estimate the maximum tumor size of our synthetic tumors in physical units. To do so, we treat our one-dimensional simulations as projections along a spherically growing tumor. We assume a typical radius of cancer cells of $r_{\text{cell}} \approx 10\,\mu\text{m}$. This allows us to estimate the volume of one LGCA voxel via its capacity $K$, assuming that at the carrying capacity, the voxel is fully occupied with cells. Therefore $V_{\text{node}} = 100 V_{\text{cell}} = 100 \cdot \frac{4}{3}\pi r_{\text{cell}}^3 \approx 4.2 \cdot 10^5\,\mu\text{m}^3$. Thus, we estimate the side length of one voxel as $L_{\text{node}} \approx 75\,\mu\text{m}$. Since the lattice is almost fully filled at the end of our simulations and we start with cells in the middle of the lattice, the final tumor diameter is approximately $100 \cdot 75\,\mu\text{m} = 0.75\,\text{cm}$.

#### 3.2 Recurrence time

To estimate the time step length in real units, we consider the cell cycle time of glioblastoma cells *in vitro*. For these cells, effective cell cycle times (neglecting births of non-viable daughter cells) $T_b = 38 \pm 4\,\text{h}$ were reported [1]. This value should correspond to that of a single cell in our model, i. e., one that is not inhibited in its growth by other cells. The cell cycle time and growth rate $\alpha$ are connected using the time step length $\tau$ via

$$\alpha(\tau) = \frac{\tau}{T_b} \Leftrightarrow \tau = \alpha T_b. \tag{S18}$$

For $\alpha = 1$ we have $\tau = T_b \approx 38\,\text{h} \approx 1.6\,\text{d}$.

### References


1. Hegedüs B, Czirók A, Fazekas I, Bábel T, Madarász E, and Vicsek T. Locomotion and proliferation of glioblastoma cells in vitro: statistical evaluation of videomicroscopic observations. J Neurosurg. 2000; 92:428–34. DOI: 10/dqxds8
